# Are seasonally plastic anti-predatory and desiccation tolerance traits developmentally linked?

**DOI:** 10.64898/2026.05.19.726136

**Authors:** Bhanu Bhakta Sharma, Ullasa Kodandaramaiah

## Abstract

In many tropical areas, seasonal rainfall leads to distinct dry and wet seasons. Many butterflies developing under wet season conditions develop into adults with large ventral eyespots on the wing margins, whereas those developing under dry season conditions have smaller or no eyespots. In greener, wet season habitats, larger eyespots can divert predator attacks toward the wing margins, while reduced eyespot size improves camouflage in the dry leaf litter-dominated habitat during the dry season. However, the dry season is also characterised by higher desiccation stress than the wet season. We hypothesised that larvae developing under dry season conditions develop into adults with higher desiccation tolerance than those reared under wet season conditions. We tested this by rearing larvae of the butterfly *Mycalesis mineus* under simulated dry and wet season conditions and assaying the desiccation tolerance of the resulting adults. Butterflies reared in dry conditions survived longer under desiccation stress, lost lesser water during pupal–adult metamorphosis, and were heavier than those reared in wet conditions. We also tested the correlation between eyespot size and desiccation tolerance. A negative correlation between the traits would be expected if similar developmental pathways regulate them. Consistent with this expectation, individuals with smaller eyespots had higher desiccation tolerance. Our results demonstrate plasticity in desiccation tolerance, and suggest that predator avoidance and desiccation tolerance traits may share similar developmental pathways.

## INTRODUCTION

In most tropical areas, seasonal change in rainfall leads to distinct dry and wet seasons (Feng et al., 2013; S. Halali et al., 2021; Kemp, 2001; Kuricheva et al., 2021). Dry seasons are characterised by low relative humidity (RH) and wet seasons by high RH (Benoit, 2010; de Souza et al., 2025). Seasonal shifts in moisture impose several physiological challenges on terrestrial organisms. During the dry season, insects are particularly vulnerable to desiccation due to their small body size which results in high surface area-to-volume ratio, relatively high metabolic rates, and proportionately low fat storage (reviewed in Bujan et al., 2016; Gibbs et al., 1997; O’Donnell, 2022). To cope with desiccation stress, insects have evolved a suite of traits that can reduce water loss, increase tolerance to water loss, or enhance water storage capacity (reviewed in Gibbs et al., 2003; Parkash & Aggarwal, 2012; Sinclair et al., 2024). For instance, some insects modulate their metabolism by lowering metabolic rates or reallocating metabolites to different physiological functions (Chown et al., 2011; Hoffmann & Parsons 1989; Marron et al., 2003; Matzkin & Markow, 2009), while other species avoid desiccation stress by entering aestivation, an extreme form of dormancy in which the metabolic rate is drastically reduced (Benoit, 2010; Huestis & Lehmann, 2014). Behavioural responses, such as seeking shaded, moist microhabitats and reducing activity during the driest hours of the day are also widespread strategies (Benoit et al., 2023; Bujan et al., 2016; Kessler & Guerin, 2008). Because much of water loss is through the cuticles (Gibbs & Rajpurohit, 2010; Kühsel et al., 2017; Rourke, 2000), desiccation tolerance in many insects involves modifications of cuticular properties including increased melanin pigmentation (Farnesi et al., 2017; King & Sinclair, 2015; Rajpurohit et al., 2008; Ramniwas et al., 2013) and deposition of longer-chain hydrocarbons (Gibbs, 1998).

Because desiccation tolerance traits can be resource-intensive, expressing them during wet seasons might be counterproductive (Chown et al., 2011; Kwan et al., 2008). For example, low metabolic rate conserves water in dry conditions but can be disadvantageous in wet conditions, where rapid growth and reproduction are favoured (Hoffmann & Parsons, 1989). This illustrates a classic ‘resistance life-history trait trade-off’, wherein traits that enhance survival under stressful conditions often reduce performance under favourable conditions (Alpert, 2006; Bujan et al., 2016; Holmes & Benoit, 2019). Phenotypic plasticity — the ability of a genotype to produce different phenotypes depending on the developing environment (West-Eberhard, 1989; Whitman & Agrawal, 2009) allows insects to express desiccation tolerance traits only when beneficial. For e.g., desiccation tolerance traits differ across seasonal populations in *Drosophila* species (Kalra & Parkash, 2016; Rajpurohit et al., 2018). Thus, insects are able to sense changes in the environment, use cues to anticipate the future environment and selectively express desiccation tolerance related traits only during the dry season (reviewed in Aggarwal et al., 2013; Bosua et al., 2023; Kellermann et al., 2018; Sinclair et al., 2024).

Seasons also differ in terms of selection imposed by predators. Wing pattern polyphenism in butterflies is an oft cited example of adaptive phenotypic plasticity that allows species to cope with seasonal changes in predation pressure (Bhardwaj et al., 2020; Brakefield & Reitsma, 1991; Molleman et al., 2024; Pfennig, 2021). Such polyphenism has been reported in many tropical butterflies and particularly well studied in *Bicyclus anynana* (Beldade et al., 2002; Lyytinen et al., 2004; van Bergen & Beldade, 2019; van Bergen & Oostra, 2023; Windig et al., 1994). Seasonal forms differ in eyespot size: wet season forms have large, prominent ventral eyespots along the wing margins, whereas these eyespots are reduced or absent in dry season forms (Braby, 1994; Brakefield & Larsen, 1984). These wing patterns serve distinct functions. In the wet season, the marginally located eyespots deflect predator attacks away from vital body parts (Halali et al., 2019; Prudic et al., 2015). In contrast, reduced eyespots of the dry season morph improve camouflage against the dry season background dominated by dry leaf litter (Halali et al., 2019; Lyytinen et al., 2004; Prudic et al., 2015).

In many tropical regions temperature strongly predicts seasons, with dry seasons being cooler than wet seasons (Brakefield & Reitsma, 1991). In such regions, butterflies can use temperature as a cue to develop seasonal forms (Brakefield & Larsen, 1984; Windig et al., 1994). Studies on multiple butterflies have shown that larvae reared at lower and higher temperatures develop into adults of the dry and wet season morph, respectively (Braby, 1994; Brakefield & Larsen, 1984; Kooi & Brakefield, 1999; Mallick et al., 2024; Mateus et al., 2014; Molleman et al., 2024; van Bergen et al., 2017). On the other hand, temperature is a poor predictor of season in many regions, and recent studies have shown that butterflies from such regions can use RH as cue for eyespot size plasticity (Prasannakumar et al., 2025; Yumnam et al., 2025).

In addition to predation, seasonal forms are also likely to be differentially selected by climatic factors, including moisture availability. Thus, survival in the dry season likely also requires a strategy to minimize water loss and tolerate high desiccation stress. While the adaptive significance of wing pattern polyphenism is well characterised (Lyytinen et al., 2004; Prudic et al., 2015), little is known about the plasticity of desiccation tolerance in response to seasonal climatic changes or its potential correlation with wing pattern plasticity. We here investigate desiccation tolerance of dry and wet season forms, and explore hypotheses related to the link between desiccation tolerance and wing pattern.

We reared larvae of a satyrine butterfly, *Mycalesis mineus* (Linnaeus, 1758), at low and high RH, and assayed desiccation tolerance of adults developing from these larvae. We predicted that, under desiccation stress, butterflies reared at low RH show reduced water loss compared to butterflies reared at high RH. We measured water loss during metamorphosis, i.e., during the transformation from pupa to adult, because this is a critical developmental stage where substantial water is lost (Chown & Nicolson, 2004; Tsukamoto et al., 1987), and water loss at this stage can strongly affect survival success (discussed in Childs, 2015; Rajpurohit et al., 2021). We also predicted that, under desiccating conditions, butterflies reared at low RH survive longer than those reared at high RH. Furthermore, we posited that the plasticity of eyespots and desiccation tolerance-related traits are developmentally linked. This could be because the environmental cues or developmental pathways regulating the two types of plasticity are shared. Because eyespots are smaller in the dry season than in the wet seasons, eyespot size can be used to quantify wing pattern forms (Brakefield & Reitsma, 1991; de Jong et al., 2010; Mallick et al., 2024; Molleman et al., 2024; Prasannakumar et al., 2025). Thus, we predicted that eyespot size is inversely correlated to survival probability under desiccation stress. Because large body size is associated with higher desiccation tolerance in insects (Guedes et al., 2015; Kühsel et al., 2017; Lighton et al., 1994), we also predicted that body size will be higher at low RH.

## MATERIALS AND METHODS

### Butterfly rearing

*Mycalesis mineus* females were collected from a rubber plantation and placed in an aluminium cage (40 × 40 × 40 cm) for oviposition on potted maize plants (*Zea mays*). Ripened banana slices were provided *ad libitum* as a food source for the adults. Newly hatched larvae were transferred onto 10-14 day old maize plants, and the plants were placed individually in nylon mesh sleeves (0.135m x 0.28m x 0.95m). Each sleeve contained 20-22 larvae, and was randomly assigned to one of two treatments that differed in RH (i) Low RH: 60% RH, and representing dry conditions (ii) High RH: 85% RH, representing humid conditions. A 12:12 hour light:dark cycle and a temperature of 27 °C were used for both treatments. The growth conditions of the two treatments were achieved using growth chambers (model E-36VL; Percival Scientific Inc., IA, USA), with the treatments alternated between the two growth chamber units periodically. Vapour Pressure Deficit (VPD) in the two treatments was calculated using the formula VPD = ((100 – RH)/100)*SVP, where VPD is vapour pressure deficit and SVP is saturated vapour pressure (Bujan et al., 2016). The VPD of the 27 °C, 85% RH condition was 0.535 kPa and that of 27 °C, 60% RH was 1.426 The monthly average VPD for the study area (Supp Mat. Fig. S1) ranges from 0.439 to 1.388 kPa, where the lower value reflects the most humid month and the higher value the driest. Therefore, the growth chamber conditions used in our experiment approximate the natural extremes experienced in the field.

The positions of the experimental sleeves within the growth chamber were randomised daily. Plants were changed on alternate days. Sleeves were checked daily for pupation. Larval developmental time for each individual was recorded as the number of days from hatching to pupation. On the second day of pupation, pupae were gently separated from the substrate of attachment, and their weight (hereafter, pupal weight) recorded using a Mettler Toledo JB1603-C microbalance (Mettler-Toledo, Switzerland) with a precision of 0.001 g. Each pupa was kept in a 100 ml Tarson tube with a fold of tissue paper at the bottom and covered with nylon mesh at the top. Pupae were maintained under the same growth conditions as the larvae. Four-day-old pupae were transferred to a black container (500 ml) with tissue paper at the bottom and covered with nylon mesh at the top. The containers provided enough space for the successful eclosion of butterflies. The black container ensured dark conditions that reduced the activity of eclosed butterflies and minimized wing damage. Butterfly eclosion was monitored every day at a fixed time (around 18:00 hrs). Eclosed butterflies were separated from uneclosed individuals and exposed to desiccation after 24 hours. The 24 hour window after eclosion ensured that the butterflies released all the meconium before the desiccation assay.

### Desiccation survival assay

One day after eclosion, each butterfly was weighed to record the initial (i.e., pre-desiccation) body weight. The butterfly was placed inside a custom built transparent polythene envelope for weighing. The initial body weight is hereafter referred to as ‘fresh weight’. Butterflies were then individually introduced into 500 ml transparent plastic containers containing 8 g of the desiccant silica gel (Sigma-Aldrich, 13767) placed at the base. The desiccant was pre-dried in an oven at 60°C (Thermo Scientific, Heratherm oven) for a minimum of 12 h to remove any moisture. A single layer of tissue paper was placed above the silica gel to prevent direct contact between the butterfly and the desiccant. The containers were sealed tightly. The RH inside the container ranged between 10-20% (monitored using a HTC PDF LOG datalogger, Hazari Tech Connect, Mumbai). All containers were placed in a growth chamber (model E-36VL; Percival Scientific Inc., IA, USA), maintained at 27 °C with a 12:12 light-dark cycle.

Survival was monitored every 6 hours for the first 24 hours, and every 3 hours thereafter, until death. The container was gently shaken at each scheduled time point to assess responsiveness. Butterflies showing no movement were considered dead. Upon death, individuals were immediately removed, and their post-desiccation weight was recorded. The same microbalance was used for all weight measurements in this study. The number of hours each butterfly survived under desiccation conditions was considered its desiccation survival time (hereafter ‘survival’). Specimens were then oven-dried at 60 °C (GLAB India Digital Bacteriological Incubator, India) for a minimum of 48 hours, after which their dry weight was measured (adopted from Gibbs et al., 1997; Mazer & Appel, 2001).

Percentage and absolute water content, and metamorphosis water loss were calculated as follows:

#### Percentage water content

100 x (Fresh weight – Dry weight) / Fresh weight (Mazer & Appel, 2001)

#### Absolute water content

(Fresh weight – Dry weight) (Gibbs & Matzkin, 2001)

#### Metamorphosis water loss

100 x (Pupal weight – Fresh weight) / Pupal weight (Molleman et al., 2024). Weight loss from the pupal stage to adult eclosion is assumed to be due to water loss.

### Eyespot measurement

Butterfly wings were photographed under bright light conditions after dry weight measurements. Wings of each butterfly were positioned flat on a neutral grey card, and images were captured using a Nikon D3200 (Nikon, Tokyo, Japan) digital camera with a 105 mm fixed focal length lens (Sigma Corporation, Kanagawa, Japan). Eyespot size relative to wing span was quantified for both forewings and hindwings following established protocols (Molleman et al., 2024; Prasannakumar, Molleman, Chandavarkar, et al., 2025; van Bergen et al., 2017) in ImageJ v.1.52a (Schneider et al., 2012) using a custom macro. Briefly, three landmarks were placed on each wing, and the area within them was used as a proxy for wing area. For forewing and hindwing, the eyespot Cu1 was measured (Fig. 1). Forewing eyespot size was quantified from the diameter of the outer black ring, assuming the eyespot was circular (Yumnam et al., 2025). Since the hindwing eyespot was elliptical, its size was quantified by drawing two lines: the longest (roughly parallel to the Cu1 vein) and the shortest (roughly perpendicular to the vein and passing through the pupil, Fig. 1) (Yumnam et al., 2025). Relative eyespot size was obtained by dividing eyespot area by the proxy wing area, and used for further analyses (Prasannakumar et al., 2025; Yumnam et al., 2025).

**Figure 1.**
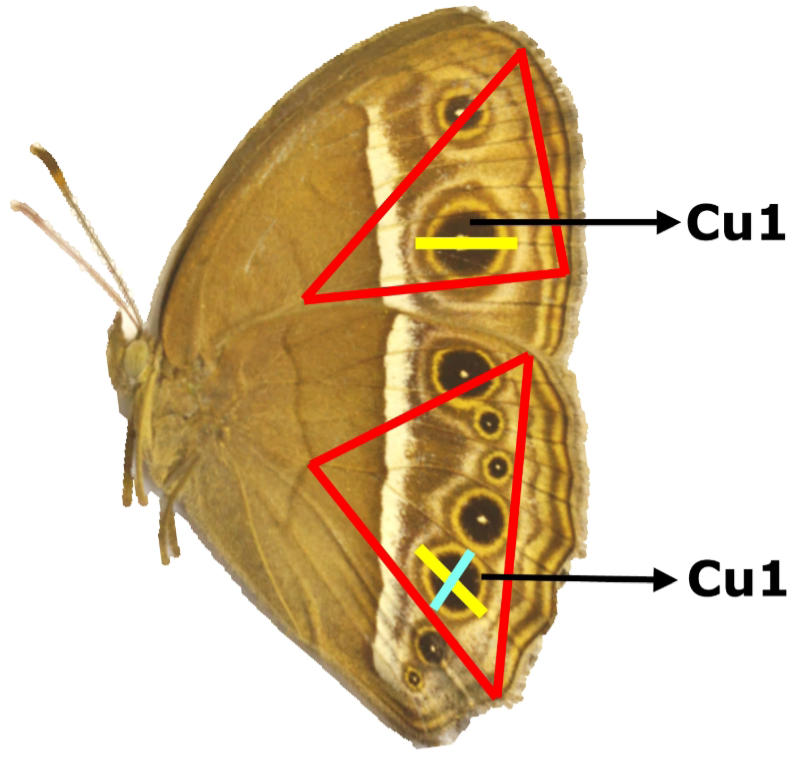
**Wing measurement landmarks of *M. mineus*** The red triangle indicates the region used to estimate wing area. For forewing yellow line marks the diameter used to measure eyespot area while for hindwing yellow line is the longest line while cyan line is the shortest.

### Statistical analyses

Data analyses and visualisation were performed in R v. 4.5.2 (R Core Team, 2024) via R Studio v. 2025.9.2.418 ‘Cucumberleaf Sunflower’ (Posit Team, 2024). Spearman correlation tests were first done to test whether forewing and hindwing eyespot sizes were correlated across individuals. The tests were done independently for each sex and experimental treatment, using the function *‘cor.test’.* Because forewing and hindwing eyespot sizes were strongly positively correlated in all four cases, ie., across both sexes and treatment (Supp. Mat. Table. S1), only forewing eyespot sizes were used for further analyses. Shapiro–Wilk tests with the *‘shapiro.test’* function indicated that the continuous variables – fresh weight, metamorphosis water loss, percentage water content, and absolute water content – all deviated from normality. Inspection of quantile–quantile plots using the function *‘descdist’* in the package *fitdistrplus* v. 1.1.14 (Marie Laure Delignette-Muller & Christophe Dutang, 2015) indicated that the gamma distribution was the best fit for all these variables.

#### Relationship between RH and eyespot size

Because relative forewing eyespot size values (0.001-0.571) range between 0 and 1, the relationship between RH and forewing eyespot size was modelled using beta regression (Douma & Weedon, 2019; Geissinger et al., 2022) using the function ‘*glmmTMB*’ in the package *glmmTMB* v. 1.1.13 (Brooks et al., 2017; McGillycuddy et al., 2025), with relative eyespot size as the response variable, and RH and Sex, and their interaction as predictor variables. As only two biological predictors were used, no model selections were performed and inference were made based on the full interaction model.

A series of Generalized Linear Models (GLMs) were built using the function *‘glm’* in the package *stats* v.4.4.2 (R Core Team, 2024) to test the effects of predictor variables on response variables (see sections *Relationship between survival and RH* and *Relationship between survival, RH, and eyespot size)*. Models were built with a gamma distribution with log link. For models with only two biological predictors, model selections was not performed and inference was made based on the full interaction model. For models with more than 2 predictor variables, subsets of the global model were obtained using the function *‘dredge’* in the package *MuMIn* v. 1.48.4 (Bartoń, 2024). Subset models were ranked based on the Akaike Information Criterion (AICc), and models having the lowest AICc values were considered the best at explaining the response outcomes (Palombo et al., 2020; Sutherland et al., 2023). Models with delta values less than 2 were averaged by using the function *‘model.avg’* in the package *MuMIn* v. 1.47.5 (Bartoń, 2024) and resulting conditional averages are reported.

#### Relationship between survival and RH

A two way GLM was built to test the combined and interactive effects of RH and sex (i.e., RH + Sex + RH X Sex) on survival. The results of the GLM analysis were corroborated with survival analyses based on log-rank tests (Group, 2004) using the function *‘survdiff’* in the package *survival* v. 3.7.0 (Therneau, 2024). The survival analyses were done independently for each sex. Kaplan-Meier survival curves were plotted independently for males and females using the function *‘ggsurvplot’* in the package *survminer* v. 0.5.0 (Kassambara et al., 2024) to visualize survival.

#### Relationship between body weight and RH

A two way GLM was built to test the combined and interactive effects of RH and sex (i.e., RH + Sex + RH X Sex) on fresh weight and dry weight.

#### Relationship between survival, RH and eyespot size

A three way global GLM first tested the combined and interactive effects of relative forewing eyespot size, RH and sex on survival. Subset models were derived and conditional averages are reported as mentioned above. The interaction between the forewing eyespot size and sex was significant. Therefore, the relationship between relative forewing eyespot size on survival was analysed independently for each sex.

#### Mediation of effect of RH on survival

Pathways linking RH to survival were estimated using structural equation models (SEM) (Britton & Davidowitz, 2024; Debecker et al., 2015) using the ‘sem’ function in the *lavaan* package v. 0.6.21 (Rosseel et al., 2012). SEM allows the simultaneous estimation of direct and indirect effects of a predictor on a response variable via multiple mediators (Gunzler et al., 2013). We built sequential mediation models. Four traits - metamorphosis water loss, percentage water content, fresh weight, and relative forewing area - were included as parallel mediators, while sex was included as a covariate. Fresh weight and dry weight were positively correlated (Supp. Matt, Table S2), similarly absolute water content and percentage water content were positively correlated (Supp. Matt, Table S3). Hence only fresh weight and percentage water content were used for the analyses. Since metamorphosis water loss was measured prior to all other adult traits and represents an earlier developmental process, it was specified as an early trait influenced by RH and Sex, and as a predictor of later adult traits (percentage water content, fresh weight, and relative forewing area). These later traits were treated as downstream mediators linking RH to survival. These traits were allowed to covary to reflect their shared physiological basis. Survival time (dependent variable) was treated as a continuous response, and models were fitted using maximum likelihood with robust standard errors (MLR) (Rosseel et al., 2012). RH and sex were treated as factors. Standardized path coefficients and standardized indirect effects are reported.

## RESULTS

### Relationship between RH and eyespot size

RH had a significant negative effect on relative forewing eyespot size (GLMM estimate = –0.374 ± 0.136, p = 0.006, Fig. 2), with butterflies reared under low RH developing smaller eyespots than those reared under high RH. The effect of sex (GLMM estimate = –0.463 ± 0.134, p = 0.005) was significant while its interaction with RH was not significant (GLMM estimate = –0.119 ± 0.0.209, p = 0.568).

**Figure 2.**
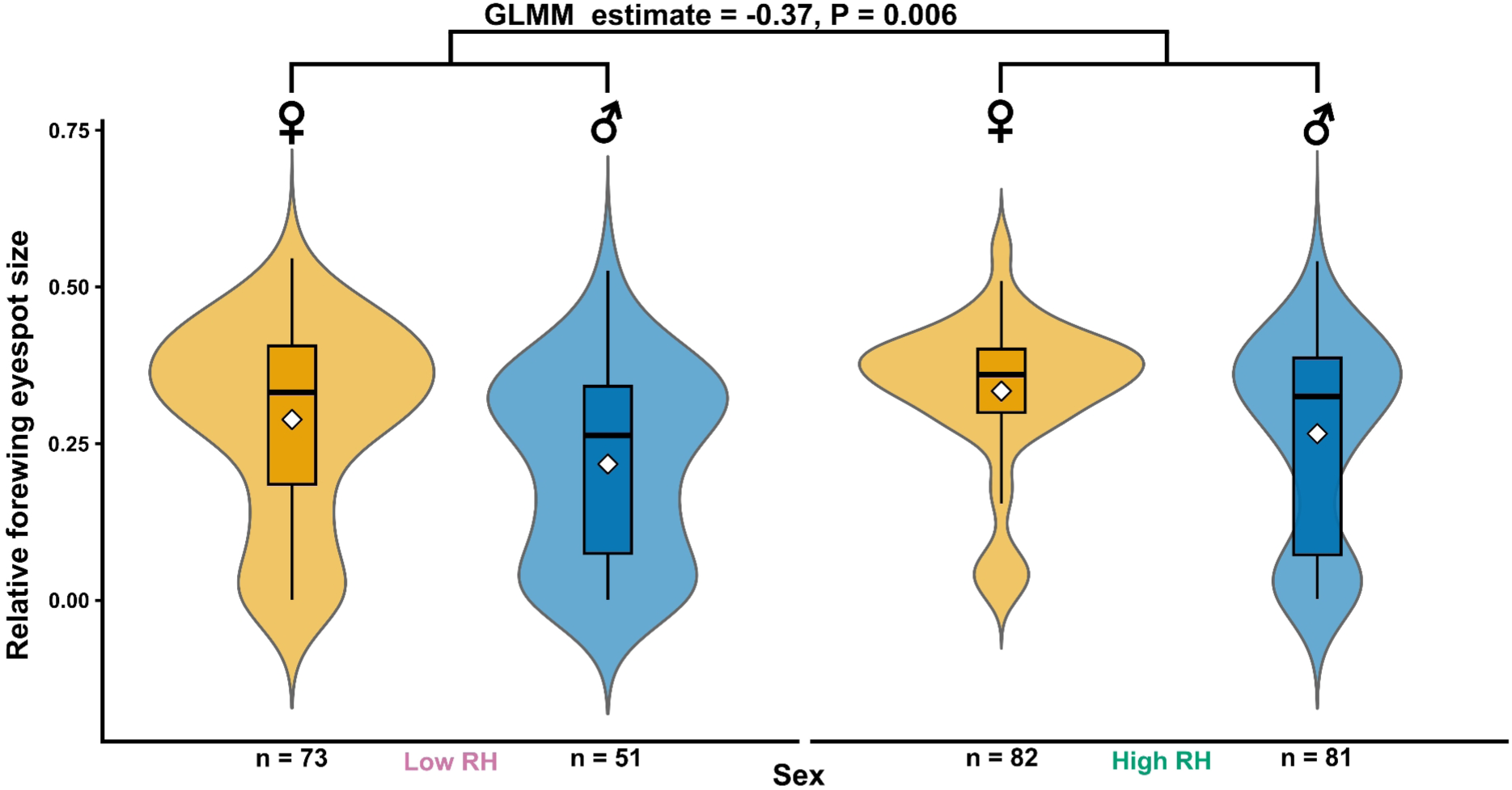
Relative forewing eyespot size across RH. Violin and box plots showing relative forewing eyespot size of individuals reared under low (left) and high RH (right). Each violin (yellow violin and box represents female while blue represents male) represents the distribution of relative forewing eyespot size with the central box showing the interquartile range (IQR), the horizontal line indicating the median, and whiskers extending to 1.5xIQR. The points beyond the whisker are outliers. The white diamond denotes the mean. Sample sizes (n) are indicated below each group. The text above the horizontal bars represent the estimate of independent effect of RH on relative forewing eyespot size derived from GLMM built to test the combined and interactive effects of RH and sex on relative forewing eyespot size.

### Relationship between survival and RH

Both GLM and log-rank survival analyses indicated that RH significantly affected survival. In the GLM, individuals reared at low RH survived longer than those at high RH (estimate = 0.106 ± 0.045, p = 0.020), and survival differ between sex (estimate = –0.108 ± 0.044, p = 0.014) while the RH X Sex interaction was not significant (estimate = 0.075 ± 0.068, p = 0.268). The log-rank test confirmed that survival differed significantly between RH treatments (χ² = 19.0, df = 1, p < 0.001), with longer survival at low RH. Independent log-rank test for each sex further showed that effect of RH was significant in both females (χ² = 7.3, df = 1, p = 0.007) and males (χ² = 10.5, df = 1, p = 0.001), with individuals of both sexes reared at low RH surviving longer than their counterparts at high RH (Fig. 3).

**Figure 3.**
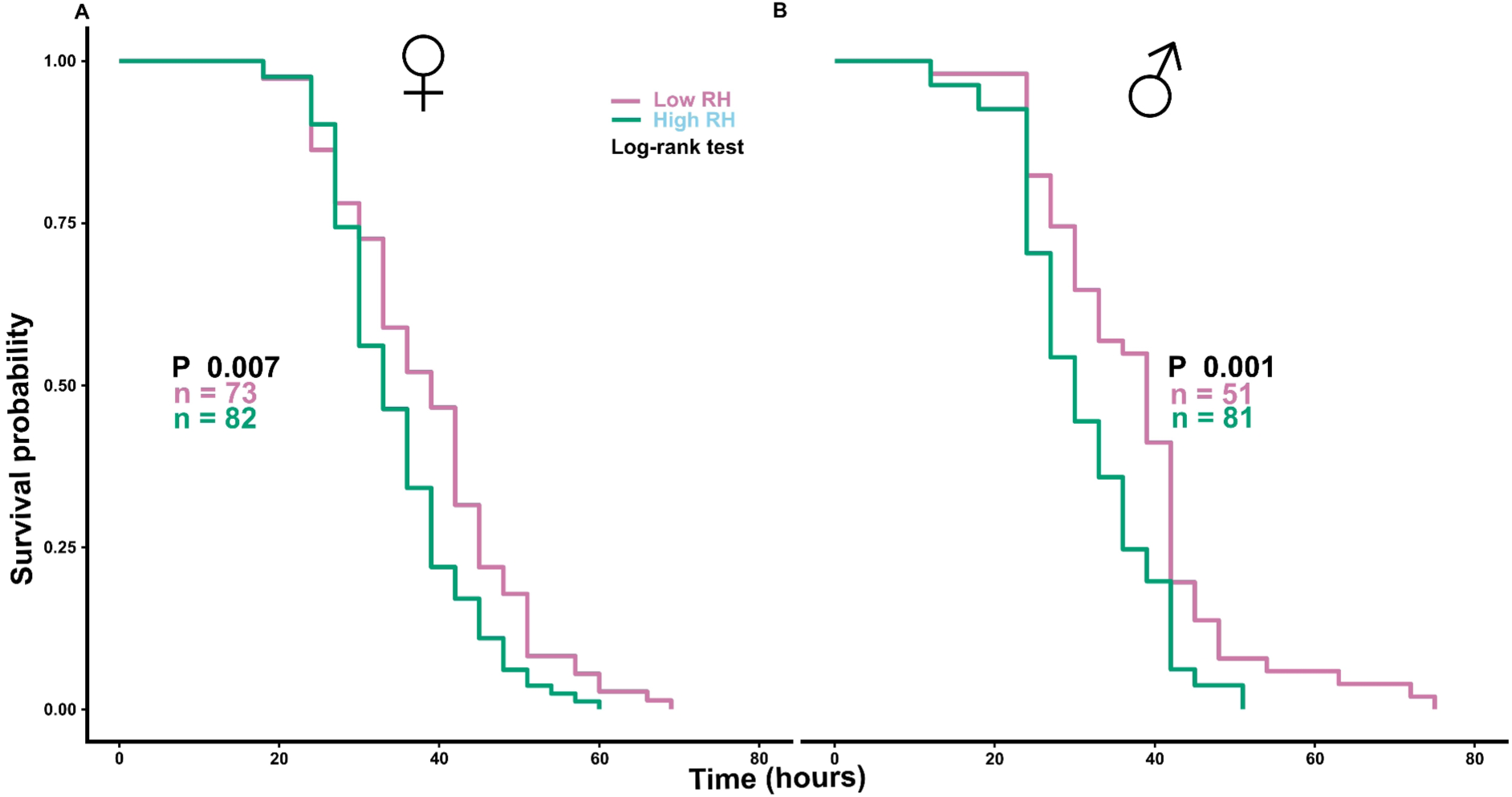
Survival probability under desiccation stress across RH. Kaplan-Meier survival curves showing survival probability of females (A) and males (B), under low (reddish purple) and high RH (bluish green). The x-axis represents the duration of survival under desiccation stress. Survival estimates were derived from independent log-rank test for each sex.

### Relationship between body weight and RH

Fresh weight differed significantly between RH treatments. Butterflies reared at low RH were heavier than those reared at high RH (estimate = 0.056 ± 0.020, p = 0.007). Fresh weight also differed between sexes (estimate = –0.319 ± 0.020, p < 0.001), with females being heavier than males. The RH X Sex interaction was not significant (estimate = 0.007 ± 0.031, p = 0.809). Similar results were obtained for dry weight: butterflies reared at low RH had higher dry weight than those reared at high RH (estimate = 0.065 ± 0.031, p = 0.036) and dry weight also differed between sexes (estimate = –0.380 ± 0.030, p < 0.001).

### Relationship between survival and eyespot size

Conditional model averaging of models including relative forewing eyespot size, sex, RH and their interaction showed that relative forewing eyespot size did not have a direct effect on survival (estimate = –0.322 ± 0.173, p = 0.06). Similarly, sex also did not have a direct effect on survival (estimate = 0.054 ± 0.07, p = 43). However, the interaction between sex and relative forewing eyespot size was significant (estimate = –0.68 ± 0.212, p = 0.001), indicating that the effect of eyespot size on survival differs between sexes (see detailed model summary in Supp. Mat. Table: S4a). Sex-specific analyses that included the interaction between relative forewing eyespot size and RH showed that relative forewing eyespot size negatively affected survival in males (estimate = –0.848 ± 0.176, p < 0.001), whereas no significant effect was detected in females (estimate = –0.43 ± 0.246, p = 0.079, Fig. 4). The independent effect of RH and its interaction with relative forewing eyespot size on survival are detailed in Supp. Mat. (Table: S4b1 and S4b1b for males and females respectively).

**Figure 4.**
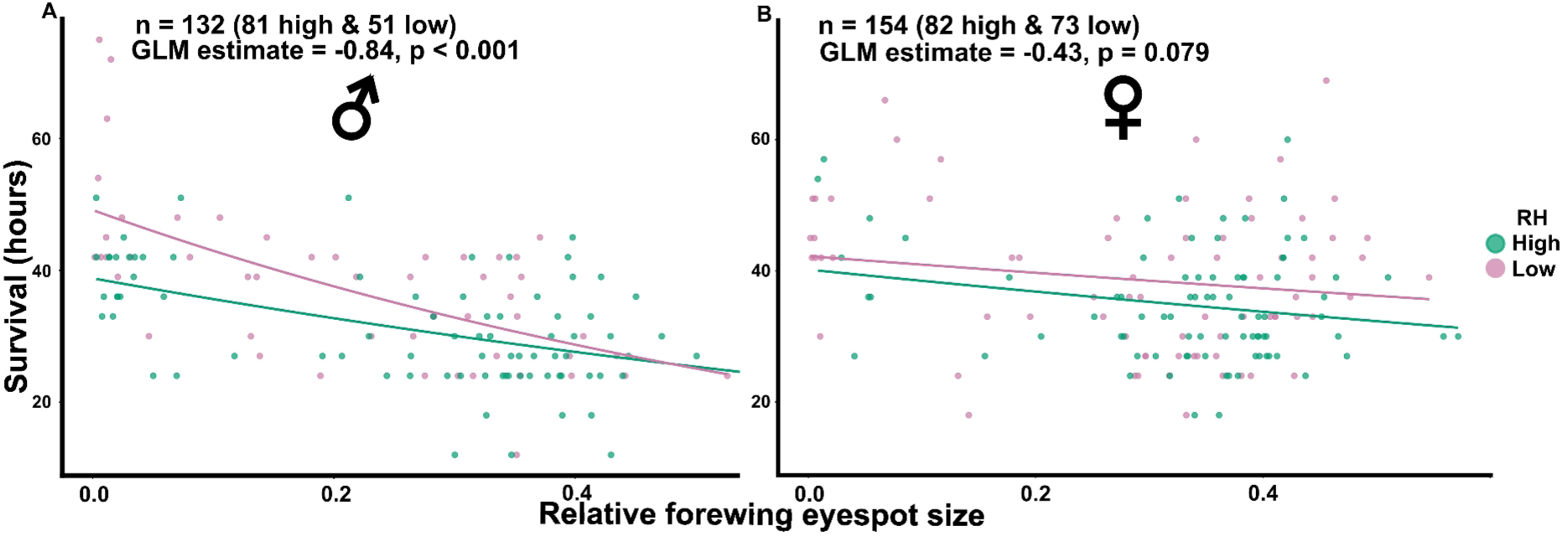
Scatter plots showing the correlation between relative forewing eyespots size and survival hours for males and females under high and low RH. Points represent individual butterflies, with RH distinguished by colour low RH (reddish purple) and high RH (bluish green). Lines on the plots show fitted trends generated using generalized linear models (gamma distribution, log link) for each sex with RH as a predictor. Estimates were derived from independent generalized linear models (Gamma distribution, log link) fitted separately for each sex in which survival was modelled as a function of the interaction between relative forewing eyespot size and RH. Panel A represents the males from low and high RH and Panel B represents the females from low and high RH.

### Pathways mediating the effect of RH on survival

The sequential SEM showed that RH influences survival both directly and through the mediators. Butterflies reared at high RH had higher metamorphosis water loss than ones reared at low RH (estimate = −2.289, z = **−**6.850, p < 0.001, Fig. 5). Metamorphosis water loss, in turn predicted two of the later adult traits: it was negatively associated with percentage water content (estimate = −0.289, z = −3.346, p = 0.001) and fresh weight (estimate = −0.526, z = −8.373, p < 0.001), but had no effect on relative forewing eyespot size (estimate = −0.013, z = −0.130, p = 897).

**Figure 5:**
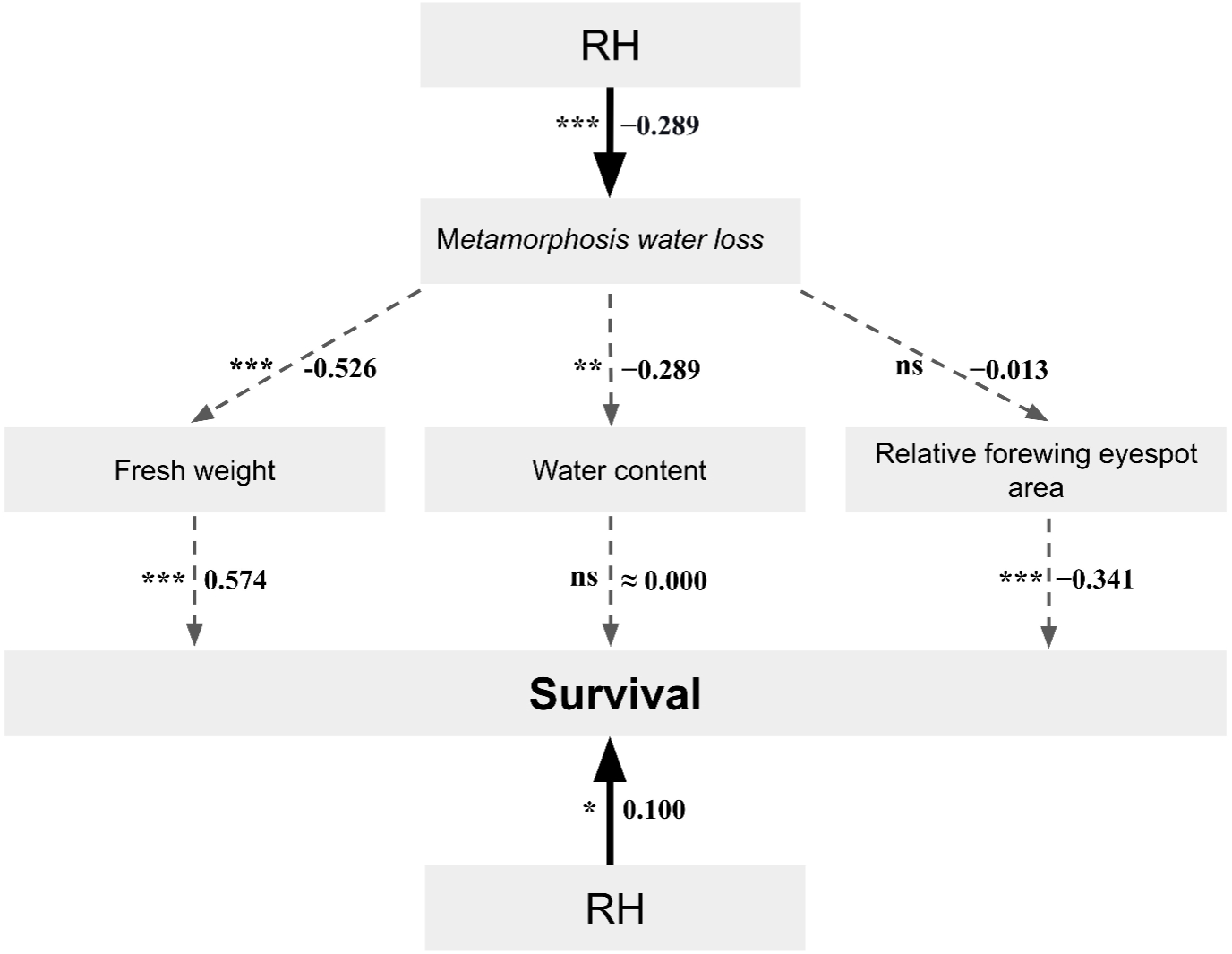
Path analysis depicting the direct and indirect pathways linking RH to survival via mediators. Numbers along paths indicate standardized path coefficients. Dashed arrows indicate indirect pathways while solid arrow indicate direct pathway. Labels “ns” indicate non significant effects, while asterisks denote levels of statistical significance (*p = 0.01-0.05, **p = 0.001-0.01, ***p < 0.001).

Survival was determined by adult traits, with positive effects of fresh weight (estimate =0.574, z = 6.794, p < 0.001) and negative effect of relative forewing eyespot size (estimate = −0.341, z = −6.357, p < 0.001). Percentage water content (estimate = −0.002, z = 0.035, p = 0.97) did not affect survival. After accounting for these pathways, RH retained a significant direct effect on survival (estimate = 0.099, z = 2.010, p = 0.044), indicating partial mediation.

Among the indirect pathways, only the route via metamorphic water loss → fresh weight → survival was significant (estimate = 0.087, z = 3.77, p < 0.001) whereas the pathways via relative forewing eyespot size (estimate = −0.001, z = −0.130, p = 0.896) and percentage water content (estimate ≈ 0.000, z = 0.035, p = 0.97) were not significant. The total (estimate = 0.186, z = 3.37, p = 0.001) and indirect effect (estimate = 0.086, z = 3.72, p < 0.001) of RH on survival was significant.

## DISCUSSION

Our results (Fig. 2) corroborate those from recent studies (Prasannakumar et al., 2025; Yumnam et al., 2025) indicating that larvae developing at low RH develop into adults with smaller eyespots. Importantly, under desiccation stress, butterflies reared at low RH survived significantly longer than those reared at high RH (Fig. 3). Mediation analysis revealed that this enhanced survival was explained by correlated mechanisms: individuals reared at low RH lost less water during metamorphosis, emerged heavier as adults, had higher fresh weight (Fig. 5). These results demonstrate adaptive developmental plasticity, wherein environmental conditions experienced during development induce expression of traits that influence adult desiccation tolerance.

An important factor determining desiccation tolerance is body weight (Kühsel et al., 2017). Large body size may confer a physiological advantage under desiccating conditions (Guedes et al., 2015; Lighton et al., 1994) because of greater absolute water reserves (Gefen et al., 2006; Gibbs et al., 1997; Guedes et al., 2015), which can be critical for survival. However, in our study, survival was not mediated by water content. Instead, individuals with larger body may have benefited from a lower surface area–to–volume ratio, which limits cuticular water loss (Addo-Bediako et al., 2001; Gibbs et al., 1997; Guedes et al., 2015). Larger body weight may also translate to higher amounts of metabolites such as trehalose (Gefen et al., 2006; Shukla et al., 2015), glycogen (Gibbs et al., 1997; Sawabe & Mogi, 1999), and lipids, which increase desiccation tolerance by acting as a source of bound water, or by serving as metabolic fuel (Chown & Nicolson, 2004; Danks, 2007; Graves et al., 1992). Such metabolites contribute to body weight (both fresh and dry weight) dry weight. Because our results suggest that the effect of RH on survival was mediated via body weight, our study provides indirect support for a metabolite-based mechanism.

In addition to body weight related effects, reduced water loss during metamorphosis contributed to the higher desiccation tolerance of the low RH group. Metamorphosis is typically associated with substantial water loss (Chown & Nicolson, 2004; Molleman et al., 2011; Tsukamoto et al., 1987), but individuals developing under low RH may adjust their physiology — for example, by reducing cuticular water loss or metabolic water loss — to minimise this loss. (Chown & Nicolson, 2004). By conserving water during this stage, adults emerging under dry season conditions may retain sufficient reserves, thereby increasing their tolerance to desiccation.

Eyespot size was inversely correlated with survival under desiccation stress (Fig. 4). Thus, during the dry season, the cryptic dry morph is better protected not only against predation (Halali et al., 2019; Prudic et al., 2015) but also against the desiccation stress typical of this season. The wet morph employs deflective eyespots to survive better against predation during the wet season (Halali et al., 2019; Prudic et al., 2015), and also invests fewer resources into desiccation-related traits, potentially allowing greater investment into other traits that increase fitness. Thus, selection by both predation and desiccation likely drives the evolution of seasonal polyphenism in butterflies.

RH may independently influence predation and desiccation-related traits through distinct developmental pathways (van Bergen et al., 2017). Alternatively, both sets of traits may represent components of adaptive developmental plasticity coordinated by shared physiological regulators such as hormones (Mateus et al., 2014). In this study, there was a correlation between eyespot size and survival not only across, but also within the RH treatments, although the correlation within treatments appears to be weak in females (Fig. 3). Thus, even among males developing at the same RH, individuals with more extreme dry forms survived better than those with wet forms. This correlation suggests that eyespot expression and desiccation tolerance may be developmentally linked, although further studies are needed to confirm the underlying mechanisms.

## SUMMARY AND CONCLUSIONS

Our findings demonstrate that desiccation tolerance is developmentally plastic in butterflies. Individuals reared under low RH survived longer under desiccation stress. This increased survival appears to be mediated by better water conservation and greater body weight. The inverse correlation between eyespot size and desiccation tolerance suggests a functional link between eyespots and stress tolerance, with cryptic morphs favoured in dry seasons and conspicuous morphs in wet seasons. These results highlight developmental plasticity as an adaptive strategy that enables butterflies to cope with contrasting seasonal conditions. Further studies are needed to ascertain whether the inverse relationship between eyespot size and desiccation tolerance reflects a shared developmental regulation, and to understand how multiple selective forces interact to shape adaptive strategies in fluctuating environments.

### Data Accessibility Statement

Upon acceptance of this manuscript, all data and analysis code used in this study will be archived in a public GitHub repository.

