## Supplementary file for "Are seasonally plastic anti-predatory and desiccation tolerance traits developmentally linked?"

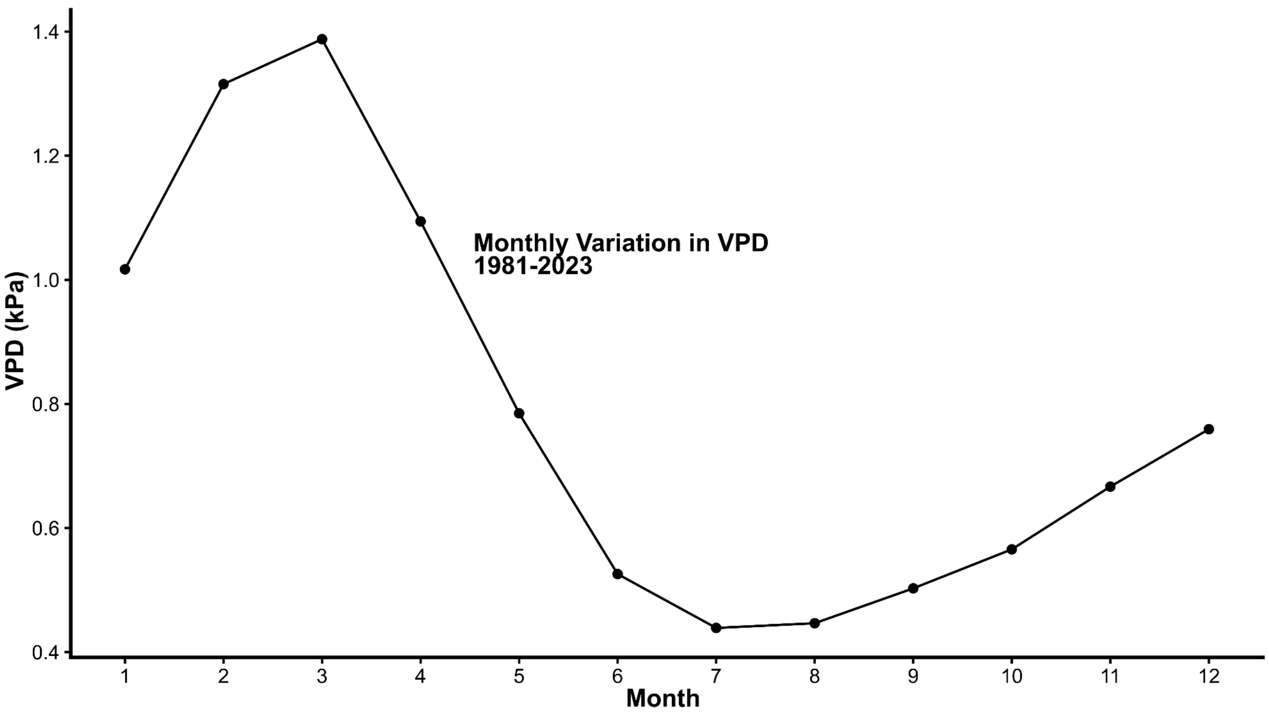


**Figure S1:** Monthly vapour pressure deficit (VPD, kPa) calculated from climate data (1981-2023) obtained from the NASA POWER database (https://power.larc.nasa.gov/data-access-viewer) for the study area. Mean monthly temperature (°C) and Relative Humidity (%) were first averaged for each month and these monthly means were then used to compute VPD using the equation VPD = ((100 – RH)/100) X SVP (Bujan et al., 2016). SVP is saturation vapour pressure and was calculated by using the Tetens formula (Wang et al., 2025).

------------------------------------------------------------------------------------------------------------------------

**Table S1**: Group wise Spearman correlations between relative forewing eyespot area and relative hindwing eyespot area, computed separately for each RH X Sex combination. For each group, Spearman's ρ, the associated p value, and the sample size (n) are reported.

| RH | Sex | Rho (ρ) | p | n |
|---|---|---|---|---|
| Low | Female | 0.792 | < 0.001 | 73 |
| Low | Male | 0.776 | < 0.001 | 51 |
| High | Female | 0.549 | < 0.001 | 82 |
| High | Male | 0.746 | < 0.001 | 81 |

------------------------------------------------------------------------------------------------------------------------

**Table S2:** Group wise Spearman correlations between fresh weight and dry weight, computed separately for each RH × Sex combination. For each group, Spearman's ρ, the associated p value, and the sample size (n) are reported.

| RH | Sex | Rho (ρ) | p | n |
|---|---|---|---|---|
| Low | Female | 0.622 | < 0.001 | 73 |
| Low | Male | 0.291 | 0.038 | 51 |
| High | Female | 0.836 | < 0.001 | 82 |
| High | Male | 0.579 | < 0.001 | 81 |

------------------------------------------------------------------------------------------------------------------------------------------

**Table S3**: Group wise Spearman correlations between absolute water content and percentage water content, computed separately for each RH × Sex combination. For each group, Spearman's ρ, the associated p value, and the sample size (n) are reported.

| RH | Sex | Rho (ρ) | p | n |
|---|---|---|---|---|
| Low | Female | 0.662 | < 0.001 | 73 |
| Low | Male | 0.868 | < 0.001 | 51 |
| High | Female | 0.393 | < 0.001 | 82 |
| High | Male | 0.609 | < 0.001 | 81 |

------------------------------------------------------------------------------------------------------------------------------------------

**Table S4a**: GLM with Gamma distribution (log link) was fitted, with survival modelled as a function of relative forewing eyespot area, RH, sex, and their interactions. Candidate models were then generated and ranked using AICc. Models with ΔAICc < 2 were retained and combined and estimates are reported.

| Variable | Estimate | Std. Error | t values | Pr(>\|t\|) |
|---|---|---|---|---|
| Intercept | 3.651 | 0.061 | 58.80 | < 0.001 |
| Relative forewing eyespot area | -0.322 | 0.173 | 1.857 | 0.063 |
| RHlow | 0.116 | 0.053 | 2.175 | 0.029 |
| SexM | 0.054 | 0.07 | 0.776 | 0.437 |
| Relative forewing eyespot area X SexM | -0.689 | 0.212 | 3.229 | 0.001 |
| Relative forewing eyespot area X RHlow | -0.195 | 0.210 | 0.921 | 0.357 |
| RHlowXSexM | 0.031 | 0.062 | 0.498 | 0.618 |

------------------------------------------------------------------------------------------------------------------------------------------

**Table S4b**: Sex-specific Gamma GLMs was fitted, with survival as a function of relative forewing eyespot area, RH, and their interaction.

***Table S4b1a** for Males*

| Variable | Estimate | Std. Error | t values | Pr(>\|t\|) |
|---|---|---|---|---|
| Intercept | 3.658 | 0.054 | 67.23 | < 0.001 |
| Relative forewing eyespot area | -0.848 | 0.176 | -4.810 | < 0.001 |
| RHlow | 0.236 | 0.082 | 2.873 | 0.004 |
| Relative forewing eyespot area X RHlow | -0.493 | 0.294 | -1.679 | 0.095 |

***Table S4b1b** for Females*

| Variable | Estimate | Std. Error | t values | Pr(>\|t\|) |
|---|---|---|---|---|
| Intercept | 3.693 | 0.087 | 42.17 | < 0.001 |
| Relative forewing eyespot area | -0.435 | 0.246 | -1.763 | 0.079 |
| RHlow | 0.049 | 0.110 | 0.448 | 0.654 |
| Relative forewing eyespot area X RHlow | 0.127 | 0.321 | 0.395 | 0.693 |

---------------------------------------------------------------------------------------------------------------------------------------
